# Ensemble of deep learning language models to support the creation of living systematic reviews for the COVID-19 literature

**DOI:** 10.1101/2023.01.18.524571

**Authors:** Julien Knafou, Quentin Haas, Nikolay Borissov, Michel Counotte, Nicola Low, Hira Imeri, Aziz Mert Ipekci, Diana Buitrago-Garcia, Leonie Heron, Poorya Amini, Douglas Teodoro

## Abstract

**Background:** The COVID-19 pandemic has led to an unprecedented amount of scientific publications, growing at a pace never seen before. Multiple living systematic reviews have been developed to assist professionals with up-to-date and trustworthy health information, but it is increasingly challenging for systematic reviewers to keep up with the evidence in electronic databases. We aimed to investigate deep learning-based machine learning algorithms to classify COVID-19 related publications to help scale-up the epidemiological curation process.

**Methods:** In this retrospective study, five different pre-trained deep learning-based language models were fine-tuned on a dataset of 6,365 publications manually classified into two classes, three subclasses and 22 sub-subclasses relevant for epidemiological triage purposes. In a *k*-fold cross-validation setting, each standalone model was assessed on a classification task and compared against an ensemble, which takes the standalone model predictions as input and uses different strategies to infer the optimal article class. A ranking task was also considered, in which the model outputs a ranked list of sub-subclasses associated with the article.

**Results:** The ensemble model significantly outperformed the standalone classifiers, achieving a F1-score of 89.2 at the class level of the classification task. The difference between the standalone and ensemble models increases at the sub-subclass level, where the ensemble reaches a micro F1-score of 70% against 67% for the best performing standalone model. For the ranking task, the ensemble obtained the highest recall@3, with a performance of 89%. Using an unanimity voting rule, the ensemble can provide predictions with higher confidence on a subset of the data, achieving detection of original papers with a F1-score up to 97% on a subset of 80% of the collection instead of 93% on the whole dataset.

**Conclusion:** This study shows the potential of using deep learning language models to perform triage of COVID-19 references efficiently and support epidemiological curation and review. The ensemble consistently and significantly outperforms any standalone model. Fine-tuning the voting strategy thresholds is an interesting alternative to annotate a subset with higher predictive confidence.

## Background

The pandemic coronavirus disease 2019 (COVID-19), caused by severe acute respiratory syndrome coronavirus 2 (SARS-CoV-2), has led to a historic wave of scientific publications in the biomedical literature (1,2). As of the beginning of the pandemic, scientific publications related to SARS-CoV-2 and COVID-19 came from the most diverse domains and became available in a myriad of digital repositories (preprint servers, technical reports, peer-reviewed scientific journals, etc.) (3). This outbreak of publications grew at an unprecedented rate. In this context, it became challenging for medical experts and epidemiologists to follow the latest scientific developments, and for curators to manually review and annotate all the available COVID-19 literature to consolidate the fast-moving existing body of knowledge (1).

Several methods for producing living systematic reviews have been proposed to provide up to date support for professionals dealing with the pace, amount and complexity of the COVID-19-related literature (4–7). A living systematic review describes a review methodology that allows updating information as soon as new evidence becomes available, rather than the methods applied to classic, time-restricted systematic reviews (8,9). Moreover, living evidence can narrow the gap between knowledge and practice, as fresh publication findings are swiftly integrated in scientifically informed guidelines (5,6,9). However, the maintenance of living evidence systems still require continuous manual curation from highly qualified human resources (10,11). One of the most time-consuming tasks is to screen the titles and/or abstracts resulting from a literature search and to exclude articles that are clearly ineligible, which may comprise a third or more of all records (2).

To address this paradigm, (semi-)automatic curation systems based on text mining and natural language processing (NLP) technologies have been developed to support review and annotation of large literature corpora (12–22). These systems support the identification and ranking of relevant articles, the categorization of the selected documents in classes and subclasses for reviewing procedures, and enable information extraction from text passages (e.g., identification of disease passages). For example, Textpresso Central (16) provides a platform that allows users to create a customized annotated corpus by uploading and processing documents of their choosing. Once documents are loaded, personalized curation searches and pipelines can be applied. PubTator Central (19) is a service for viewing and retrieving bioconcept annotations in full text biomedical articles. It comprises state-of-the-art text mining models for annotation of several biomedical entities, such as genes and proteins, diseases, chemicals, and species. SIBiLS (20) provide an optimized search engine in the biological literature by augmenting its contents with keywords and standardized entities. Variomes (22) is a system that can perform triage of publication to support evidence-based decision. Finally, PubTerm (13) enables the organization of abstracts by terms, using the co-occurrence of terms or by specific phrases, among others, to facilitate the biomedical curation process.

Automatic text classification appears as an essential methodology to ensure high quality of living evidence updates. Text classification consists of assigning categorical labels to a given text passage (e.g., an abstract) based on its similarity to the existing labeled examples (23–25). Classical text classifiers use statistical document representations, in which the relevance of a word to a document is proportional to its frequency in the document and inversely proportional to its frequency in the collection (the so-called term frequency-inverse document frequency (tf-idf) framework), to create a vectorial representations of the documents (26). These representations are then used in machine learning models, such as logistic regression and k-nearest neighbors, to learn a mapping function between the input text and the output classes (27,28). The trained models can then predict the predefined labels for new input representations. These models are however limited as they essentially fail to capture the sequential nature of text and the context in which words are embedded.

To overcome the limitations of the tf-idf framework, state-of-the-art text classifiers use deep learning-based language models to create word and document contextual representations, with improved syntactic and semantic features (29). Language models are a particular type of probabilistic model that, given a sequence of words, compute the probability distribution of the next word. Recent deep learning-based language models, such as the Bidirectional Encoder Representations of Transformers (BERT) (30), learn word representations considering both the forward- and backward-direction contexts of a word using a masked word approach, in which random words are masked from a context and the algorithm tries to predict the most likely hidden word. The models are then trained on large corpora, resulting in better word and document representations. These representations are further used as input to other NLP tasks, including text classification and question-answering, in a process called transfer learning, which has resulted in significant improvements of the state-of-the-art performance in the past years (31).

In this article, we investigated the use of automatic text classifiers supported by deep learning-based language models to enhance literature triage and annotation in COVID-19 living systematic review systems. Our analysis assessed the effectiveness of different individual deep learning-based language classifiers against two ensemble strategies, in which individual models are combined using either the probability sum of the predictions or a voting strategy where each classifier has a voting right and the classification decision is given to the class obtaining a majority of votes (32–34).

## Methodology

### Study design

An overview of the study design is presented in Figure 1. In this retrospective machine learning-based study, we evaluated the performance of different deep learning text classifiers to categorize COVID-19 literature according to their publication type in the COVID-19 Open Access Project (COAP) Living Evidence database aggregator, which includes publications about SARS-CoV-2 and COVID-19 from Pubmed, EMBASE, medRxiv and bioRxiv (4). Five individual classifiers were trained with the publication title, abstract and source associated with annotation categories of a living systematic review knowledge base. Publication title, abstract and source were imputed to the original dataset whenever missing. Remaining publications without title or abstract were excluded from the training and evaluation sets. Then, at inference time, the classifiers were applied to individual records to predict the publication category as output. Two ensemble strategies were created using these predictions (32,34). The first strategy uses a voting system that takes each classifier output as a vote for a class, while the second considers the sum of the class probabilities attributed by the individual classifiers. For the voting strategy, different cut-offs for the minimal number of votes were applied to compute the final class associated with the publication.

**Figure 1.**
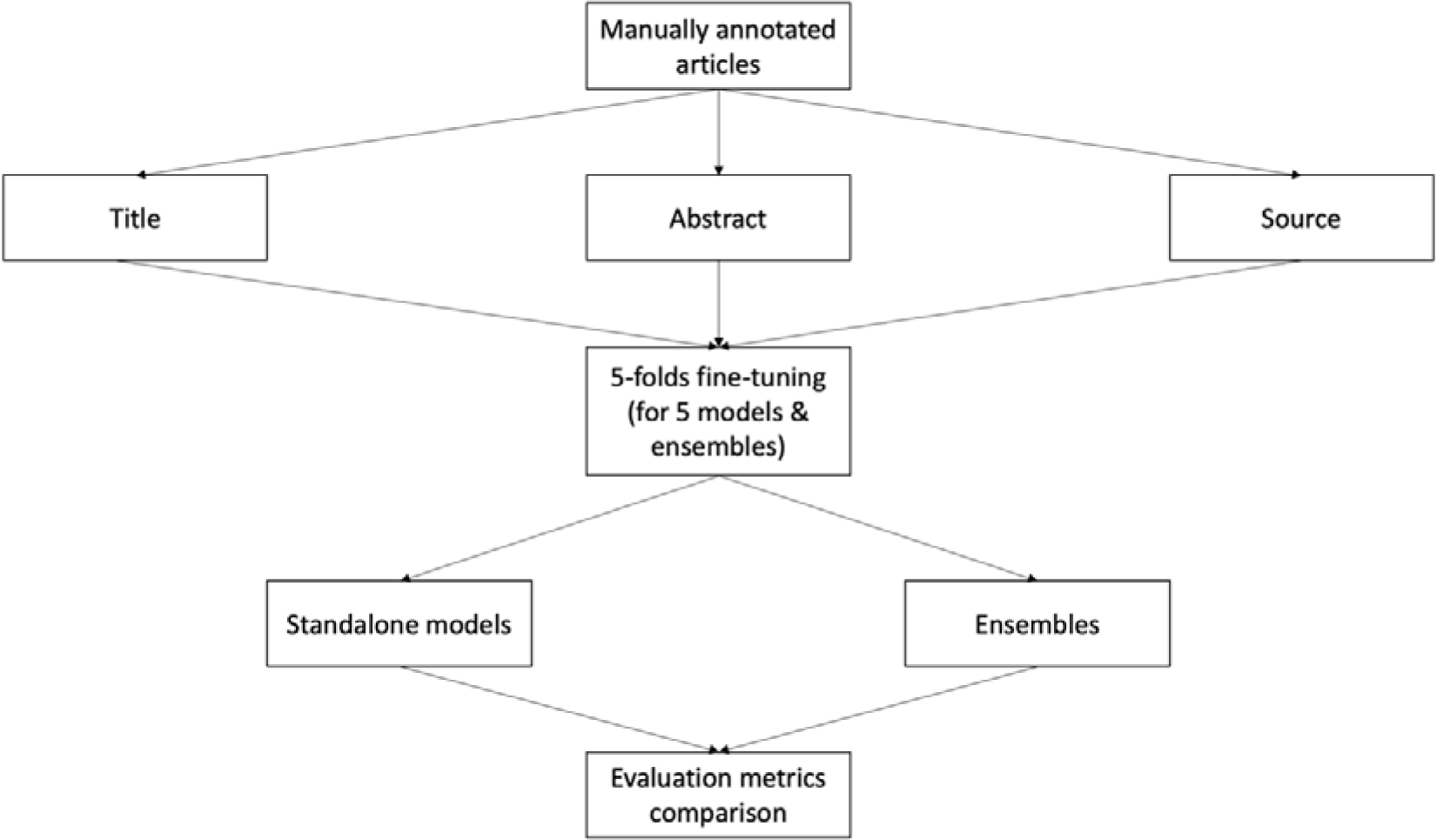
Overview of the study design. All articles were manually annotated, then the title, abstract and source retrieved. In a k-fold cross-validation setting (k is set to 5 in our experiments), 5 models were fine-tuned and each standalone model was compared against each other as well as against two types of ensemble.

Model training and evaluation were performed on a dataset of articles, which were annotated manually by a crowdsourced team of people with training in epidemiology and systematic reviews (2). Each article was manually classified across 22 sub-subclasses describing the type of COVID-19 publications according to their study design or article type (case report, ecological study, modeling study, editorial, etc.). The sub-subclasses are nested into three subclasses, namely epidemiologic study designs (EPI), basic biological or other laboratory-based research studies (BASIC) and other types of articles (OTHER). The subclasses are nested into two classes of original research (ORIGINAL) and articles that were commentaries, editorials or narrative literature reviews (NON-ORIGINAL). The source dataset is publicly available at https://zika.ispm.unibe.ch/assets/data/pub/search_beta/. To improve the robustness of the results, we trained and evaluated our models using a k-fold cross-validation methodology (k is set to 5 in our experiments). For each fold, 70% of the articles (∼4.6k publications) were used to train the model parameters, 10% unseen documents (dev set) were used to optimize the model hyperparameters and the remaining 20% unseen documents (test set) were used to evaluate the performance of the classifier. The final performance was obtained by averaging the results obtained on the *k* unseen test sets. We used standard classification metrics - precision, recall, F1-score and area under the receiver operating characteristics curve (AUC-ROC) - to assess performance of the individual models in comparison to the ensemble, and the performance of the latter at different vote majority levels (i.e., simple and absolute). The experiments were performed using the Python package HuggingFace on a Linux machine with a TPU (V3-8).

### Dataset description and pre-processing

The COAP data snapshot version used in our experiments contains 6,365 publications annotated between 7th January and 10th December 2020.Table 1A shows the distribution of publications across classes, subclasses and sub-subclasses in the COAP snapshot dataset. The categories are imbalanced for the three categorization levels, as is typically the case for real-world data. Illustratively, the *BASIC: Within-host modelling* sub-subclass composes only 0.5% of the collection (31 documents) while the *OTHER: Comment, editorial, …, non-original* sub-subclass is responsible for 27.6% (1,758 documents). There are 799 documents for the *BASIC* subclass and 3,665 documents for the *EPI* subclass, which accounts for 57.6% of the dataset. At the class level, the *ORIGINAL* class is responsible for 70.1% of the dataset, with the remaining documents (29.9%) being categorized according to the *NON-ORIGINAL* class.

**Table 1.** A: Dataset document count and proportion by class, subclass and sub-subclass. B: pre-trained models used in the experiments, the corpus type used in their training, and the number of parameters per model.

| Class | Subclass | Sub-subclass | Count | % |
| --- | --- | --- | --- | --- |
| ORIGINAL | EPI | Case report | 241 | 3.8 |
| ORIGINAL | EPI | Case series | 350 | 5.5 |
| ORIGINAL | EPI | Case-control study | 74 | 1.2 |
| ORIGINAL | EPI | Cohort study | 246 | 3.9 |
| ORIGINAL | EPI | Cross-sectional study | 284 | 4.5 |
| ORIGINAL | EPI | Diagnostic study | 181 | 2.8 |
| ORIGINAL | EPI | Ecological study | 92 | 1.4 |
| ORIGINAL | EPI | Guidelines | 326 | 5.1 |
| ORIGINAL | EPI | Modeling study | 808 | 12.7 |
| ORIGINAL | EPI | Other | 130 | 2.0 |
| ORIGINAL | EPI | Outbreak or surveillance report | 133 | 2.1 |
| ORIGINAL | EPI | Qualitative study | 35 | 0.5 |
| ORIGINAL | EPI | Review | 725 | 11.4 |
| ORIGINAL | EPI | Trial | 40 | 0.6 |
| ORIGINAL | BASIC | Animal experiment | 43 | 0.7 |
| ORIGINAL | BASIC | Basic research review | 135 | 2.1 |
| ORIGINAL | BASIC | Biochemical/protein structure studies | 264 | 4.1 |
| ORIGINAL | BASIC | In vitro experiment | 85 | 1.3 |
| ORIGINAL | BASIC | Sequencing and phylogenetics | 241 | 3.8 |
| ORIGINAL | BASIC | Within-host modeling | 31 | 0.5 |
| NON-ORIGINAL | OTHER | Other | 143 | 2.2 |
| NON-ORIGINAL | OTHER | Comment, editorial, ..., non-original | 1758 | 27.6 |
|  |  | ALL | 6365 | 100.0 |

Table 1
| Pre-trained models | Corpus type | # Parameters<br>(M) |
| --- | --- | --- |
| RoBERTa<sub>base</sub> (39) | General | 110 |
| RoBERTa<sub>large</sub> (39) | General | 340 |
| COVID-Twitter-BERT (40) | Bio (COVID-19) | 110 |
| BioBERT (41) | Bio | 110 |
| PubMedBERT (42) | Bio | 110 |

In the pre-processing phase, the title, abstract and source fields were concatenated before being fed to a classifier, and each classification model used its own tokenizer in order to separate the free text passages into tokens (words or sub-words) (35–38). All model tokenizer specificities are given in their respective papers (see Table 1B).

### Classification models

In our experiments, we used the pre-trained models shown in Table 1B, which were originally pre-trained using the masked language model task. In a masked language model task, large corpora, such as Medline or Wikipedia, are used to create low-dimensional word (or sub-words) representations in a context. In each training step, a sentence taken from the corpus is provided to the model with (sub-)words masked. The model is then trained to predict the masked (sub-)words for that context. The resulting model encodes contextualized (sub-)words in a low-dimensional space and optimal tensorial representations can then be used in downstream tasks, such as text classification, a process called transfer learning. Two out of the five models (RoBERTa-base and RoBERTa-large) were pre-trained on a general corpus, created using Book Corpus and Wikipedia, while three other models (COVID-Twitter-BERT, BioBERT and PubMedBERT) were pre-trained on biomedical corpora. Among the models trained on biomedical corpora, one was pre-trained on a COVID-19 related corpus and one can be considered as large, gathering 340M parameters. All specificities of the models can be found in their related literature (see Table 1B).

#### Individual deep learning-based classifier for biomedical literature classification

Transformer models (43) with a fully connected perceptron layer on top of the output attention layer were used to discriminate sub-subclasses of given documents. Using the pre-trained language model classifiers, knowledge acquired by the model in the pre-training phase can be transferred to the specific task, during the so called fine-tuning phase, in which task specific examples are given the original model so its parameters can be updated to the task at hand (30). In our case, the specific classification task consists of fine-tuning the models on a subset [training set] of the manually annotated dataset, followed by the classification of documents from another unseen subset [test set] among the 22 sub-subclasses of the knowledge base. At the inference phase, the model extracts features from the document metadata (i.e., title, abstract and source) and outputs a probability for each of the 22 sub-subclasses. As sub-subclasses are mutually exclusive, for a given document, the sum of all the probabilities across sub-subclasses is equal to 1. Additionally, predictions with respect to the subclass and class levels were computed. To do so, the probabilities for sub-subclasses belonging to a subclass (or classes) are summed. In other words, the probability of a document to be classified in a given class is the sum of the probabilities for that document to be classified in all the sub-subclasses mapped to that class; mapping as per Table 1A. The predicted category, i.e., class, subclass or sub-subclass, is then defined as the highest probability across all the predicted probabilities.

Figure 2 shows the publication classification workflow. The model starts with a publication containing a title, an abstract and a source. The text contained in those three fields is concatenated and a tokenizer splits it into tokens (e.g., words or sub-words). Each token is then linked to a token id which allows the language model to look up for a vectorial representation of the said token. In our example, the word “Study” is split into the “Stu” and “#dy” sub-words. “Stu” is the token id number 51 and finds its vectorial representation in the 51th model matrix row. Once retrieved, the language model will receive its vector representation v_51_ as an input along with all the other token representations. The language model then gives the publication representation to a classifier, which outputs a probability for each sub-subclass.

**Figure 2.**
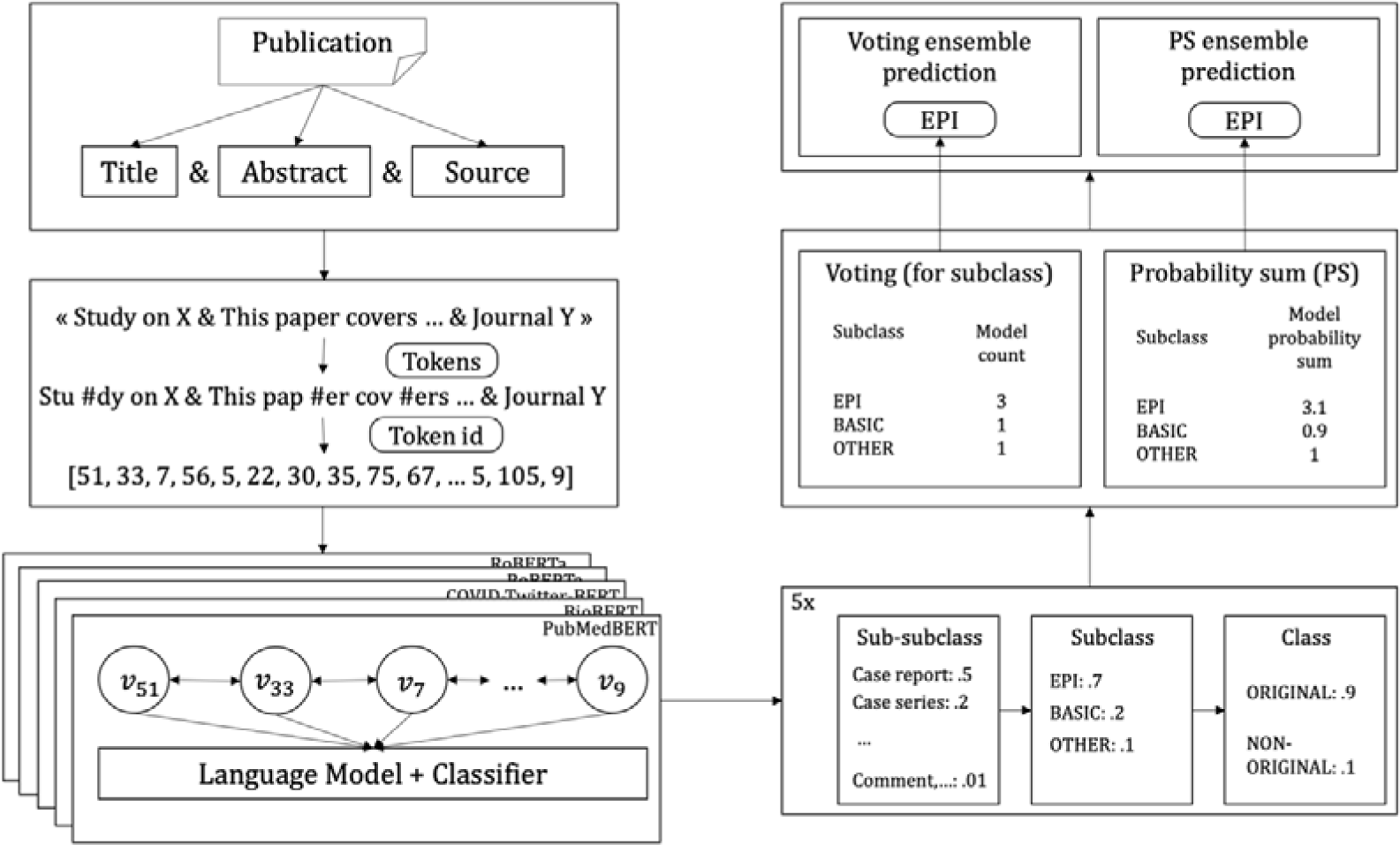
Publication classifier workflow. The model starts with the title, abstract and source fields and concatenates their text contents before tokenizing it. Each model computes their predictions and an ensemble strategy, voting or probability sum, combines them to get a final prediction.

#### Ensemble: voting and probability sum strategies

Assembling models can be performed by making individual models vote for a category. In the default version, the final category is defined by the higher number of votes. A threshold of votes which would trigger a voting ensemble prediction can also be used. In this setting, an *unknown* prediction, that is, the model is unsure about the category, is possible when there is a tie or when the number of votes is below the threshold (i.e., there is no unanimity). With this ensemble strategy, only the class level (binary) is ensured to always get predictions with a threshold equal to 3 in our setting (5 models). Alternatively, a probability sum strategy can be used to create the ensemble. The idea is to sum the probabilities of the classifiers for all the categories and then take the most probable category as the ensemble classification. If not stated otherwise, the probability sum strategy would be the default ensemble as this method always gives a unique prediction in every situation. In Figure 2, as an example, 3 out of 5 models predicted the *EPI* subclass, so the voting ensemble ended up predicting the *EPI* subclass. For the probability sum strategy, the sum of all subclass predictions among all the 5 models gives a score of 3.1 for the *EPI* subclass, which makes it the highest score among all the other subclasses. Even if in this case predictions are the same for both strategies, it is worth noting that it is not systematically the case.

### Model interpretation

To get an insight of the model word impact, the integrated gradient (44) was performed using captum (45) implementation on the PubMedBERT model on the subclass level. According to this method, the higher a token scores, the more important it is to the prediction, and the score polarity implies the positive/negative classification impact. This experiment is two-fold. First, about 600 never-seen documents were classified and the 20 highest positive impact words for each subclass prediction were reported. To deal with tokenized sub-words, a word score was computed using the mean of all its sub-words compositions. Then, to reflect a more general impact of a given word for a subclass, each word was lemmatized and the word score is computed as the mean of the respective lemmatized word scores. This way, a word and its plural would merge, for example, “simulation” and “simulations” would gather their scores and attribute their scores to the lemmatized word “simulation”. To avoid non-generalized high-impact-words, only words with at least 5 occurrences were considered. In the second part of this experiment, a few publication scores were analyzed. To do so, the set of analyzed documents sampling was driven by the top-20 positive words statistics.

### Statistical analysis

To evaluate our models, standard multiclass classification metrics were used, such as precision, recall, F1-score and AUC-ROC (26). Precision describes the proportion of correctly classified documents over all the documents being classified by the model to the same class:

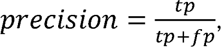

where *tp* is the number of true positives and *fp* is the number of false positives. Recall describes the proportion of correctly classified documents among all the positive documents for given class:

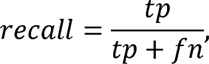

where *fn* is the number of false negatives. Finally, F1-score can be formulated as the harmonic mean of the model precision and recall:

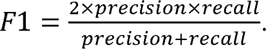

For these three metrics, the closer the result is to 1, the better is the model performance. Lastly, AUC-ROC, computes the area under ROC, where the ROC plots the curve given a classification threshold of the *tp* rate (or recall or sensitivity) against the *fp* rate (or 1 – specificity):

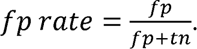

To get a confidence interval (CI) of the AUC-ROC, a bootstrapping with a sample of n=2000 was computed. The 2.5% and 97.5% values of the distribution were reported to get a 95% CI. The McNemar test is used for statistical significance testing (46).

In the ranking experiments, the model predicts a ranked list of sub-subclasses according to their probabilities for a given input document. Thus, we use standard information retrieval metrics to report our results. The precision at ranking *k* (@k) is the precision across all the first *k* sub-subclasses returned by our classifiers. As it is a multi-class problem, each document belongs only to one true class; thus, the theoretical maximum precision is equal to 1/*k*. By analogy, recall@k is set across the first *k* sub-subclasses. Conversely to precision, the more *k* increases, the more the recall@k should be close to 1. As there are 22 sub-subclasses, by definition recall@22 is equal to 1. Finally, the mean average precision (MAP) @k is the mean of all the average precisions (AP) @k, which is defined as follows:

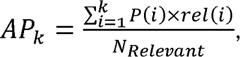

where *P(i)* is the precision at *i* position, *rel(i)* is a function equal to 1 if the *i^th^* returned document is relevant and equal to zero otherwise, and *N_Relevant_* is the number of documents relevant for a given query. As our classification problem is mutually exclusive, *N_Relevant_* is equal to 1 and *P@1 = R@1 = MAP@1*. Compared to traditional classification metrics, which only consider the top model prediction, the ranking metrics help us to understand how good are the top-k classification predictions.

## Results

### Classification performance

Table 2A-C shows the performance of the different models using the F1-score metric at the class, subclass and sub-subclass levels, respectively. The ensemble outperformed the best standalone model significantly with a micro F1-score of 89% (Table 2A). PubMedBERT obtained the best F1-score across the standalone models for all the classes. When comparing models to each other, there is no significant improvement. Although the improvement of the ensemble with respect to the PubMedBERT model is statistically significant, it accounts for less than a point for both the micro and macro F1-scores. At the subclass level (Table 2B), similarly to the class level the ensemble outperformed all single models significantly, but in this case for more than a percentage point for both micro and macro F1-scores (86% *vs.* 85% micro F1-score and 84% *vs.* 83% macro F1-score), and it is also consistently the best performing model across all the subclasses. PubMedBERT was again the overall best standalone model at the subclass level, with a micro and macro F1-scores of 85% and 83%, respectively. At sub-subclass level (Table 2C), the ensemble significantly achieved the best micro and macro average F1-score (70% and 55%), having the highest F1-score for 10 sub-subclasses, for which 3 of the improvements were statistically significant. For the standalone models, PubMedBERT had the best micro F1-score (67%), while RoBERTa*-large* presented the best macro F1-score (53%). The relevant gap between aggregated scores (micro and macro F1-scores) from Tables 2B and 2C suggests that there were more intra-level than inter-level misclassifications. In other words, misclassified sub-subclasses were often confused with sub-subclasses belonging to the same subclass. Finally, Table 2D shows the AUC-ROC performance and their respective 95% CI for each level. Here, the ensemble reports systematically a higher performance than any standalone model. When compared to BioBERT, the best standalone model in this metric, for each level there is no CI overlap, confirming the statistically significant improvement by the ensemble model.

**Table 2.**
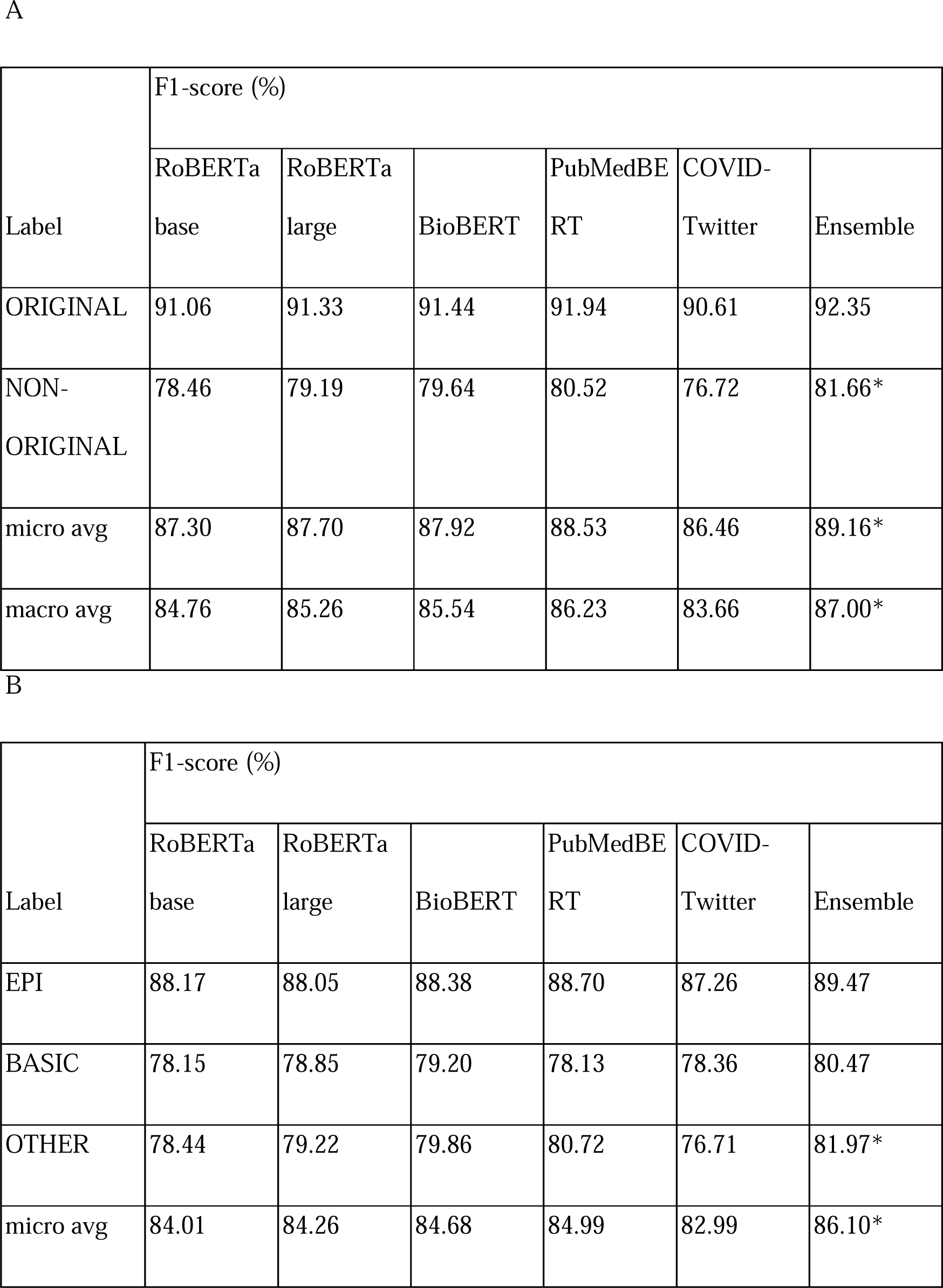
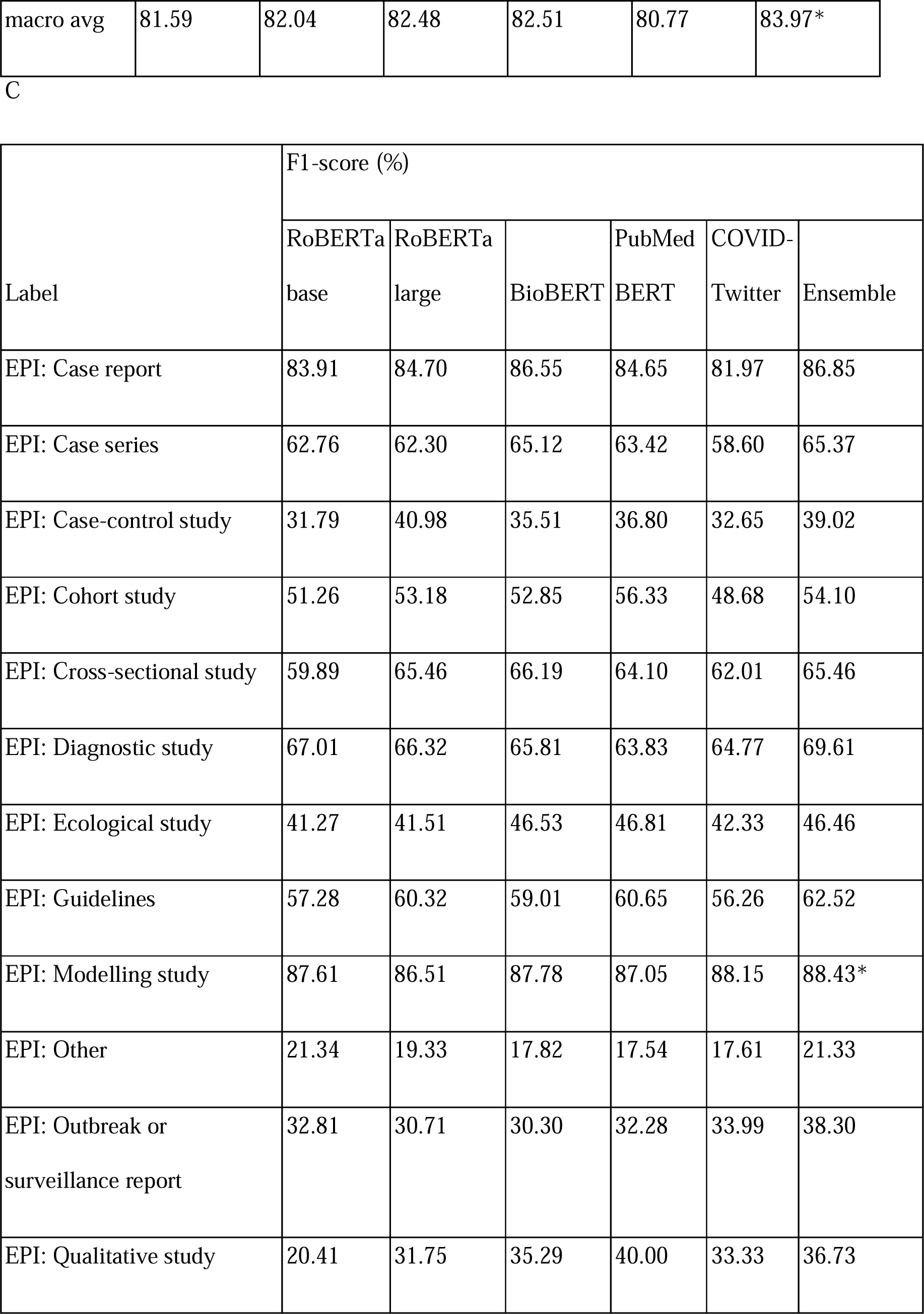
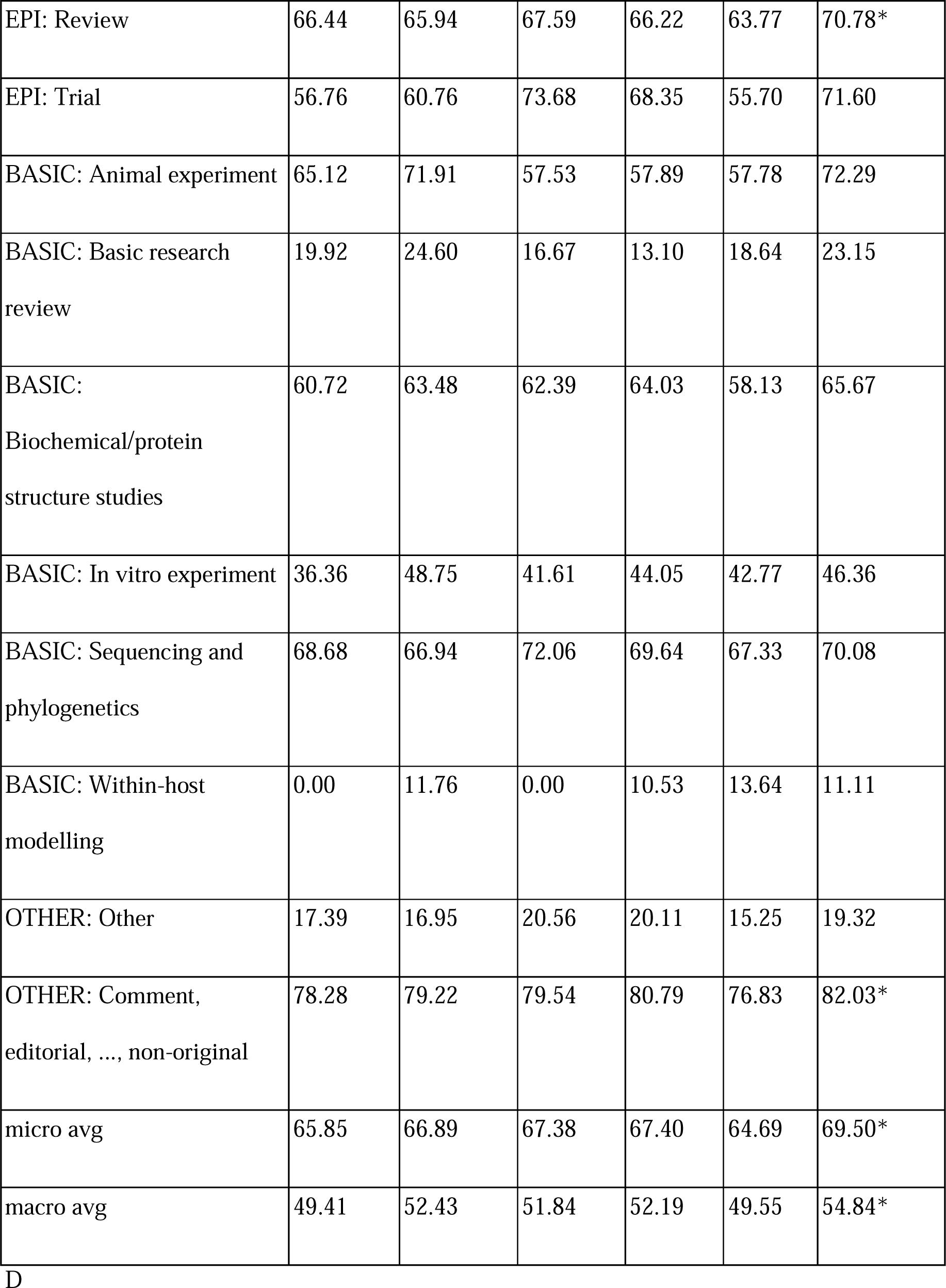
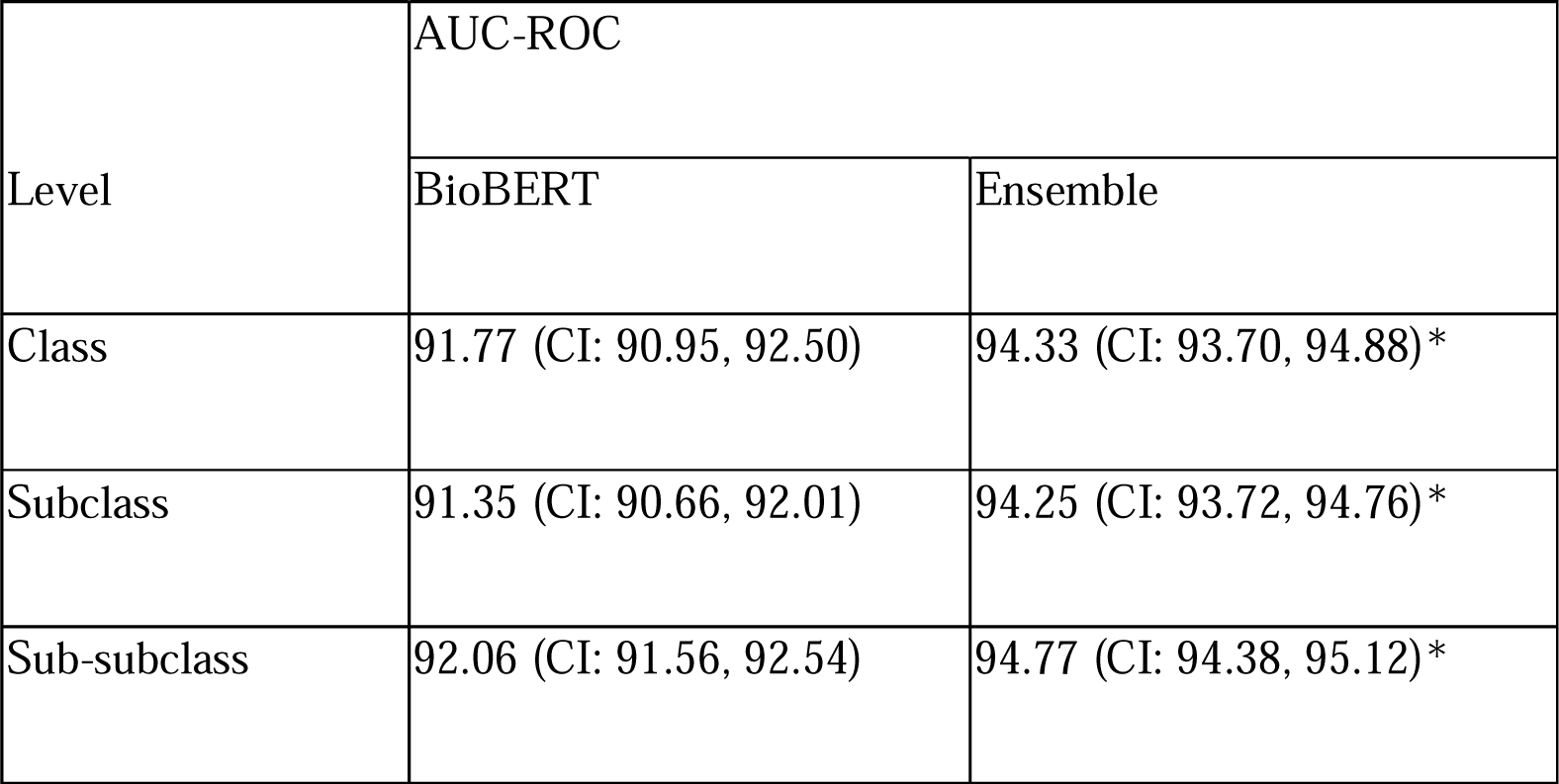
A,B,C: F1-score performance for the different models across categories and classification levels. D: AUC-ROC performance and a 95% CI for the different classification levels for the best standalone and the ensemble models. * Statistically significant improvement.

| Label | F1-score (%) |  |  |  |  |  |
| --- | --- | --- | --- | --- | --- | --- |
|  | RoBERTa<br>base | RoBERTa<br>large | BioBERT | PubMedBE<br>RT | COVID-Twitter | Ensemble |
| ORIGINAL | 91.06 | 91.33 | 91.44 | 91.94 | 90.61 | 92.35 |
| NON-ORIGINAL | 78.46 | 79.19 | 79.64 | 80.52 | 76.72 | 81.66* |
| micro avg | 87.30 | 87.70 | 87.92 | 88.53 | 86.46 | 89.16* |
| macro avg | 84.76 | 85.26 | 85.54 | 86.23 | 83.66 | 87.00* |

B
| Label | F1-score (%) |  |  |  |  |  |
| --- | --- | --- | --- | --- | --- | --- |
|  | RoBERTa<br>base | RoBERTa<br>large | BioBERT | PubMedBE<br>RT | COVID-Twitter | Ensemble |
| EPI | 88.17 | 88.05 | 88.38 | 88.70 | 87.26 | 89.47 |
| BASIC | 78.15 | 78.85 | 79.20 | 78.13 | 78.36 | 80.47 |
| OTHER | 78.44 | 79.22 | 79.86 | 80.72 | 76.71 | 81.97* |
| micro avg | 84.01 | 84.26 | 84.68 | 84.99 | 82.99 | 86.10* |
| macro avg | 81.59 | 82.04 | 82.48 | 82.51 | 80.77 | 83.97* |

C
| Label | F1-score (%) |  |  |  |  |  |
| --- | --- | --- | --- | --- | --- | --- |
|  | RoBERTa<br>base | RoBERTa<br>large | BioBERT | PubMed<br>BERT | COVID-Twitter | Ensemble |
| EPI: Case report | 83.91 | 84.70 | 86.55 | 84.65 | 81.97 | 86.85 |
| EPI: Case series | 62.76 | 62.30 | 65.12 | 63.42 | 58.60 | 65.37 |
| EPI: Case-control study | 31.79 | 40.98 | 35.51 | 36.80 | 32.65 | 39.02 |
| EPI: Cohort study | 51.26 | 53.18 | 52.85 | 56.33 | 48.68 | 54.10 |
| EPI: Cross-sectional study | 59.89 | 65.46 | 66.19 | 64.10 | 62.01 | 65.46 |
| EPI: Diagnostic study | 67.01 | 66.32 | 65.81 | 63.83 | 64.77 | 69.61 |
| EPI: Ecological study | 41.27 | 41.51 | 46.53 | 46.81 | 42.33 | 46.46 |
| EPI: Guidelines | 57.28 | 60.32 | 59.01 | 60.65 | 56.26 | 62.52 |
| EPI: Modelling study | 87.61 | 86.51 | 87.78 | 87.05 | 88.15 | 88.43* |
| EPI: Other | 21.34 | 19.33 | 17.82 | 17.54 | 17.61 | 21.33 |
| EPI: Outbreak or<br>surveillance report | 32.81 | 30.71 | 30.30 | 32.28 | 33.99 | 38.30 |
| EPI: Qualitative study | 20.41 | 31.75 | 35.29 | 40.00 | 33.33 | 36.73 |
| EPI: Review | 66.44 | 65.94 | 67.59 | 66.22 | 63.77 | 70.78* |
| EPI: Trial | 56.76 | 60.76 | 73.68 | 68.35 | 55.70 | 71.60 |
| BASIC: Animal experiment | 65.12 | 71.91 | 57.53 | 57.89 | 57.78 | 72.29 |
| BASIC: Basic research<br>review | 19.92 | 24.60 | 16.67 | 13.10 | 18.64 | 23.15 |
| BASIC:<br>Biochemical/protein<br>structure studies | 60.72 | 63.48 | 62.39 | 64.03 | 58.13 | 65.67 |
| BASIC: In vitro experiment | 36.36 | 48.75 | 41.61 | 44.05 | 42.77 | 46.36 |
| BASIC: Sequencing and<br>phylogenetics | 68.68 | 66.94 | 72.06 | 69.64 | 67.33 | 70.08 |
| BASIC: Within-host<br>modelling | 0.00 | 11.76 | 0.00 | 10.53 | 13.64 | 11.11 |
| OTHER: Other | 17.39 | 16.95 | 20.56 | 20.11 | 15.25 | 19.32 |
| OTHER: Comment,<br>editorial, ..., non-original | 78.28 | 79.22 | 79.54 | 80.79 | 76.83 | 82.03* |
| micro avg | 65.85 | 66.89 | 67.38 | 67.40 | 64.69 | 69.50* |
| macro avg | 49.41 | 52.43 | 51.84 | 52.19 | 49.55 | 54.84* |

Table 2
| Level | AUC-ROC |  |
| --- | --- | --- |
|  | BioBERT | Ensemble |
| Class | 91.77 (CI: 90.95, 92.50) | 94.33 (CI: 93.70, 94.88)* |
| Subclass | 91.35 (CI: 90.66, 92.01) | 94.25 (CI: 93.72, 94.76)* |
| Sub-subclass | 92.06 (CI: 91.56, 92.54) | 94.77 (CI: 94.38, 95.12)* |

The worst performing sub-subclasses (F1-score < 30.00), namely *EPI: Other*, *BASIC: Basic research review*, *BASIC: Within-host modelling* and *OTHER: Other*, are all underrepresented in the dataset, accounting for only 2.0%, 2.1%, 0.5% and 2.2%, respectively. The poor performance for these classes had a negative impact on the macro average F1-score, which is below the micro average for all the models. In opposition, in the best performing sub-subclasses (F1-score > 70.00), namely *EPI: Case report*, *EPI: Modelling study*, *EPI: Review*, *BASIC: Animal experiment*, *BASIC: Sequencing and Phylogenetics* and *OTHER: Comment, editorial, …, non-original,* all accounted for 3.8%, 12.7%, 11.4%, 0.7%, 3.8% and 27.6% of the dataset, respectively. Those 6 sub-subclasses (30% of the sub-subclasses) account for about 60% of the collection, yet with a high variance in their distribution. These results suggest that the number of training examples alone are not enough to explain the model performance and that textual features in the title+abstract+source fields and/or category definition make some classes easier to be learned.

### Analyses of the ensemble model

In Figure 3, we analyzed major aspects of the ensemble outcomes. In Figure 3A, the ensemble precision/recall curve is plotted against the curves for the *RoBERTa* base and large models for the *ORIGINAL* class. As we can notice, the ensemble curve is consistently above both *RoBERTa* models, which shows the robustness of using a probability sum strategy for assembling models. The precision/recall curve obtained by the ensemble model for the 22 sub-subclasses are presented in Figure 3B. The same under-performing sub-subclasses as previously spotted in the strict classification results can be distinguished, in particular, *EPI: Other*, *BASIC: Basic research review*, *BASIC: Within-host modelling* and *OTHER: Other* (as in Table 2A). This demonstrates that the low performance obtained for these categories is not a result of the classification threshold tuning. Despite their poor performance, they are well above a random classifier baseline, which would have a theoretical constant precision of about 0.05 (1/22 sub-subclasses).

**Figure 3.**
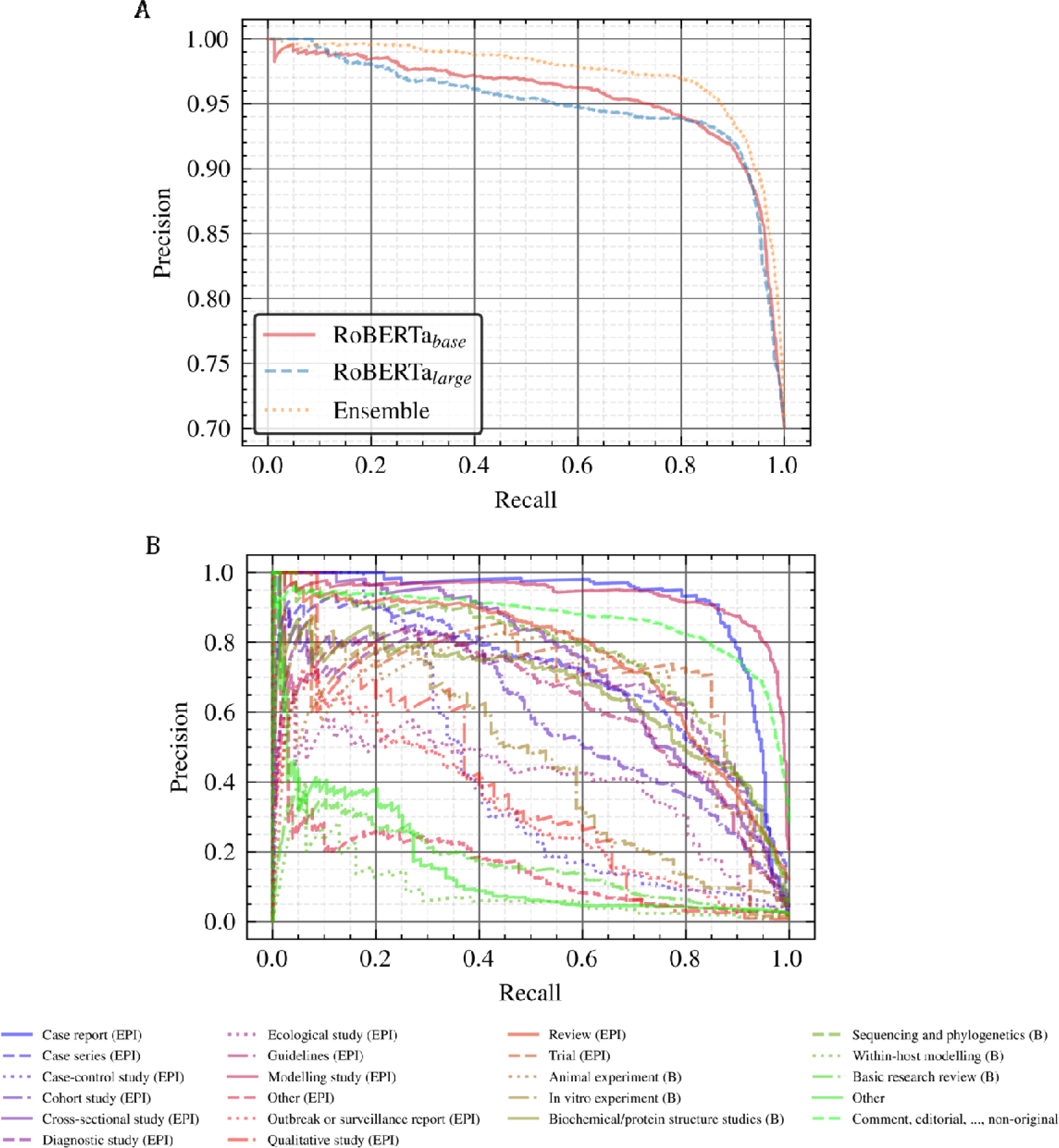
A: Precision/recall curves of the ORIGINAL class for the RoBERTa base/large and the ensemble. B: Precision/recall curves obtained by the ensemble model for the sub-subclasses. Well represented sub-subclasses usually perform better than underrepresented ones.

Figure 4 shows the confusion matrix for the different classification levels obtained by the ensemble model. As we can see from Figure 4A and 4B, the ensemble tends to predict *EPI* subclass when misclassifying a document. When switching from Figure 4A to 4B, the *EPI* confusion is split from the *BASIC* class into both *BASIC* and *OTHER*. For the sub-subclass level (Figure 4C), the *EPI: Review* class (13) was consistently confused with the *BASIC: Basic research review* (20). This confusion is expected considering that both sub-subclasses refer to review documents. Moreover, the ensemble tends to get confused for some of the *EPI: … study* sub-subclasses, predicting often *Cohort* (4) instead of *Case-control* (3), *Cross-sectional* (5) instead of *Qualitative* (12), *Modelling* (9) instead of *Ecological* (7), and others. There is also a clear confusion cluster when the ensemble predicts *Biochemical/protein structure studies* (17) and *Sequencing and phylogenetics* (18), as these documents are often confused with some of the *BASIC* sub-subclasses (in particular from 15 to 19). These observations reinforce our previous hypothesis that sub-subclasses were often misclassified inside the same subclass. It becomes more evident if we focus on the sub-subclass confusion matrix by square segments as highlighted in Figure 4C (horizontal and vertical gray lines): from index 1 to 14 ➔ *EPI*, from index 15 to 20 ➔ *BASIC* and for index 21 and 22 ➔ *OTHER*. All shady squares inside this perimeter (the majority) are intra-subclass misclassifications, while the ones outside are inter-subclass misclassifications. Lastly, a vertical line of confusion can also be observed for the *OTHER: Comment, editorial, …, non-original* sub-subclass predictions, which the ensemble tends to predict for a wide variety of documents (more precisely 8, 10-13, 20-21). The broad definition of this category is likely the reason for its confusion with so many other sub-subclasses.

**Figure 4.**
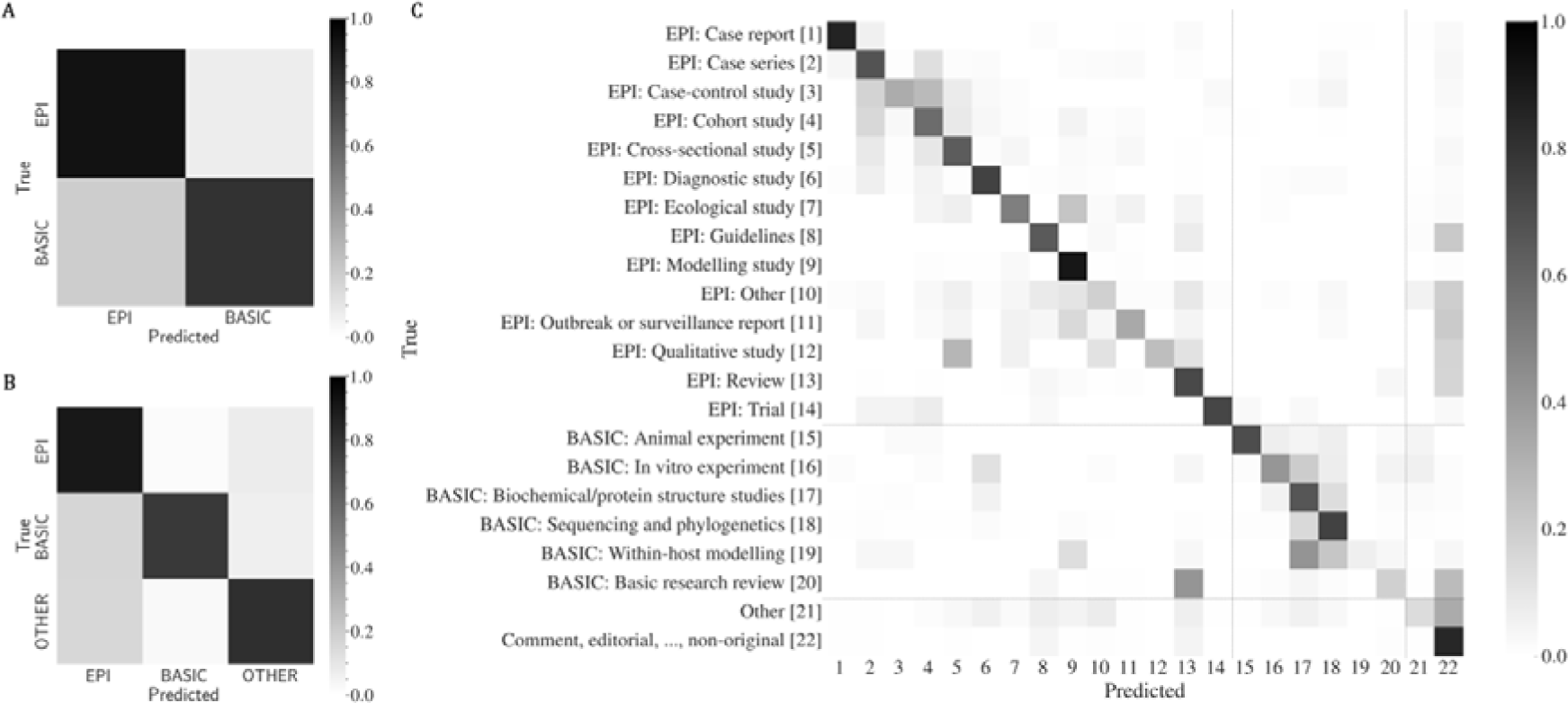
Confusion matrix for class (A), subclass (B) and sub-subclass (C). The ensemble has a higher probability of confusing sub-subclasses inside their nested subclasses and classes which is why performances tend to be higher at those higher levels.

### Ranking analysis

Table 3 shows the ranking performance for the standalone models and the ensemble. *BioBERT* performed better than all the other standalone models for the ranking metrics, whereas it tended to be *PubMedBERT* in the strict classification perspective. However, in both perspectives, the ensemble achieves the highest performance across all models. In fact, the ensemble returns the right sub-subclass in the top-1 position in 71% of cases, with precision@3 of 30% (theoretical maximum of 33%) and a recall@3 of 89%. This means that in almost 9 out 10 document classifications, the ensemble returned the correct sub-subclass in the top 3. Moreover, the ensemble got MAP@3 of 79%, representing more than 2.5 points improvement with respect to the best standalone model (*BioBERT*).

**Table 3.**
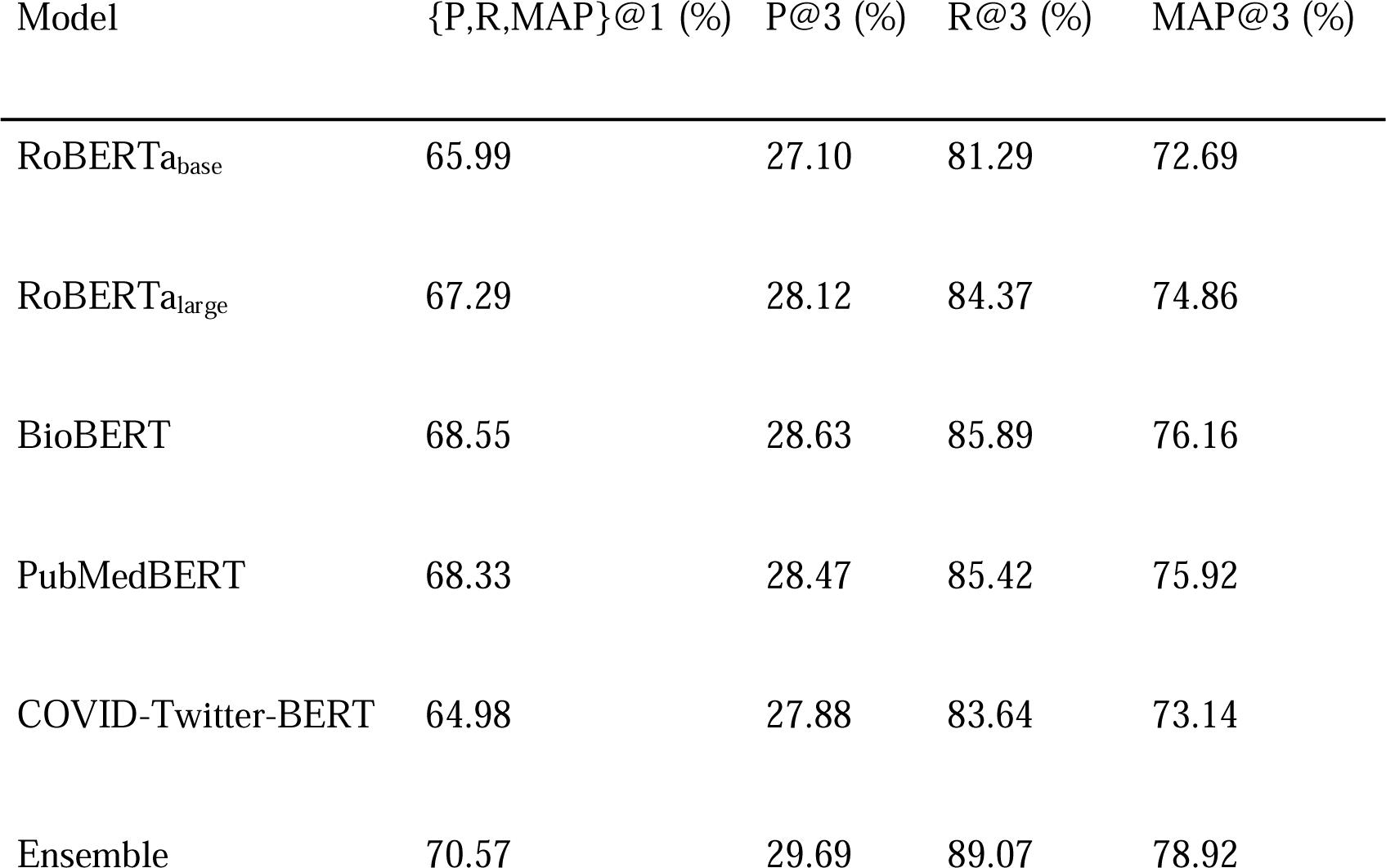
Metrics per label using the top-k retrieved categories. P=precision, R=recall, MAP=mean average precision. As this is a single-label task, the max value for P@3 is 1/3 (33%).

| Model | {P,R,MAP}@1 (%) | P@3 (%) | R@3 (%) | MAP@3 (%) |
| --- | --- | --- | --- | --- |
| RoBERTa<sub>base</sub> | 65.99 | 27.10 | 81.29 | 72.69 |
| RoBERTa<sub>large</sub> | 67.29 | 28.12 | 84.37 | 74.86 |
| BioBERT | 68.55 | 28.63 | 85.89 | 76.16 |
| PubMedBERT | 68.33 | 28.47 | 85.42 | 75.92 |
| COVID-Twitter-BERT | 64.98 | 27.88 | 83.64 | 73.14 |
| Ensemble | 70.57 | 29.69 | 89.07 | 78.92 |

### k-vote analysis

In Figures 5, we show the strict classification performance for the *ORIGINAL* class using the ensemble for different voting thresholds. The threshold for the number of votes (*t*) corresponds to the minimal number of votes for a category required for the ensemble to trigger a classification decision. Differently, the probability threshold per vote (*t_v_*) refers to the probability threshold a single model needs to reach to vote for a given category. When such a probability threshold is not met, the model would not be allowed to vote. Such voting strategies make *unknown* predictions possible, reducing the size of the classification set. In addition to static voting thresholds (3, 4, 5), a dynamic threshold, for majority and unanimity, is introduced where the total of votes can change depending on *unknown* predictions for a given classifier. This means that if 2 classifiers (out of 5) were to predict *unknown* for a publication, the dynamic majority and unanimity thresholds would be set at 2 and 3, respectively.

**Figure 5.**
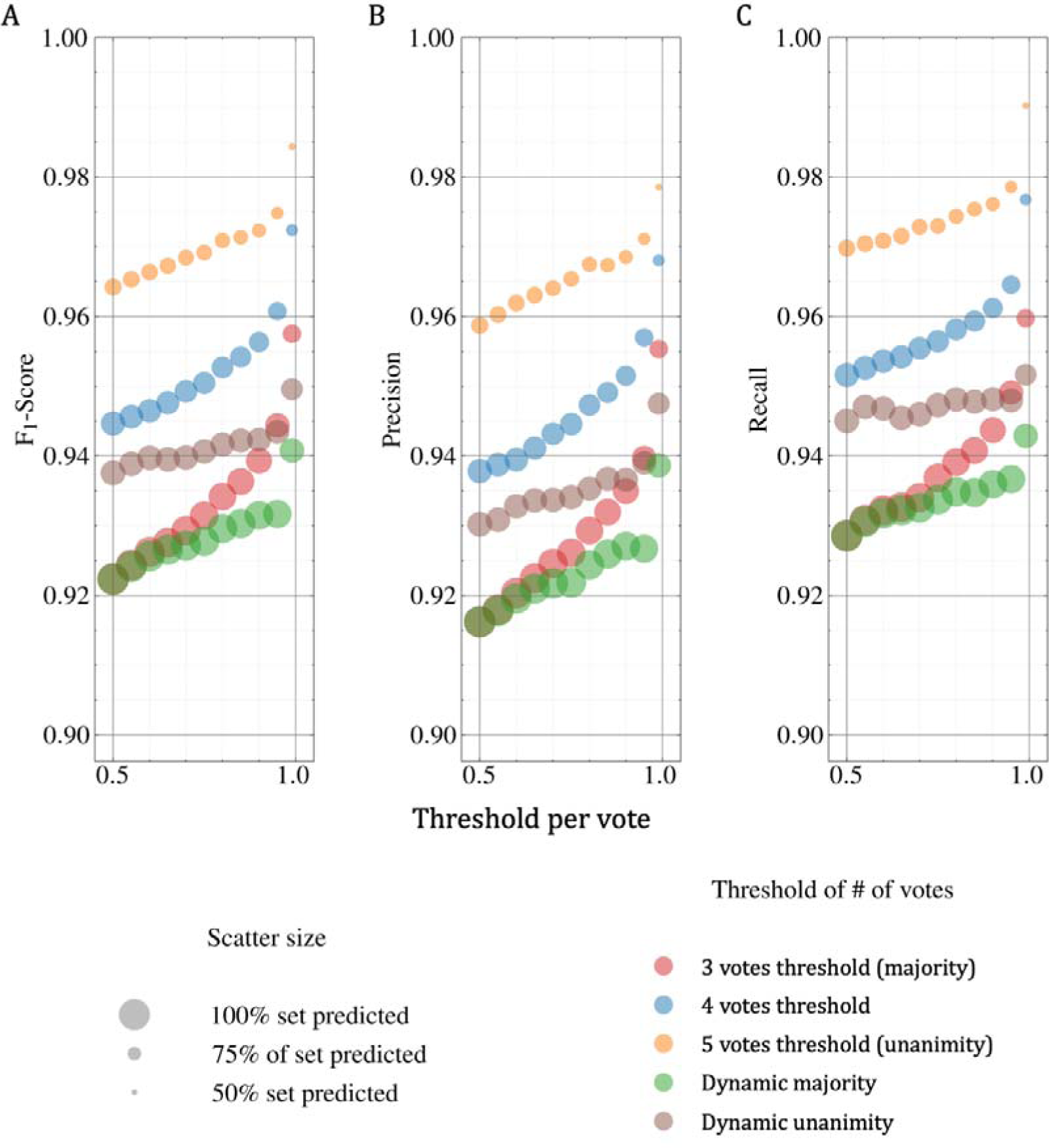
F1-score (A)/Precision (B)/Recall (C) for the ORIGINAL class with respect to a probability threshold per vote when using the voting strategy across the predictions on the class level. Using different thresholds improves considerably performance while reducing the number of predicted publications.

The behavior of the *ORIGINAL* class prediction in terms of F1-score is presented in Figure 5A. As it is a binary problem, setting a dynamic majority and a static one (*t = 3*) while *t_v_ =* 0.5 produced the same results, a full size dot placed around 92%. This phenomenon is possible because there will always be a predicted class that has more than *t_v_ =* 0.5; hence, all the models end up voting. Overall, there is an average of about 93% F1-score on most of the dataset across all the *t_v_* when using majority voting rules and 97% F1-score on a subset of about 80% of the dataset when using the static unanimity voting rule. In other words, for the *ORIGINAL* class, confident results can be obtained (about 4 points F1-score growth) on a subset of the collection (representing about 80% of the collection) when switching from a majority to static unanimity voting rule. The respective performance in terms of precision and recall metrics are shown in Figures 5B and 5C. We can notice that recall is consistently higher than precision, which means that this ensemble strategy is better at retrieving *ORIGINAL* articles than refining the selection. The observed trend is similar to the F1-score performance, where we trade a 100% dataset classification and a precision of about 91.5%, for a precision of about 96% on about 80% of the dataset with a fixed *t_v_ =* 0.5 when switching from a majority to a static unanimity voting rule. A recall of about 99% and a F1-score of about 98.5% are achieved on 50% of the subset when setting *t_v_ =* 0.99 and *t =* 5, enabling the classification of half of the publications with almost no mistakes.

### Model interpretation

Figures 6A to 6C show the top 20 positive impact words for *EPI*, *BASIC* and *OTHER* subclasses. When taking a close look at some lexical fields, in the *EPI* subclass for instance, documents containing “modeling”, “mathematical”, “modelling”, “simulation”, “simulated” and “equation” are all related to the *EPI: Modelling study* sub-subclass. Indeed, in the 38 documents subset containing at least one of those words, 37 were classified by the model as *EPI: Modelling study*. In *BASIC*, the same applies for “seq” and “sequence” lexicons, where 27 publications out of 28 were classified by the model as either *BASIC: Sequencing and phylogenetics* or *BASIC: Biochemical/protein structure studies*. In other words, the model clearly seems to retain high importance words at the sub-subclass level, which makes sense as it is the level the model was fine-tuned on. As for *OTHER*, it seems the classifier attributes a lot of credit to the word “viewpoint” for any *OTHER: Comment, editorial, …, non-original* publications, with 7 out of 7 publications containing the word classified as so.

**Figure 6.**
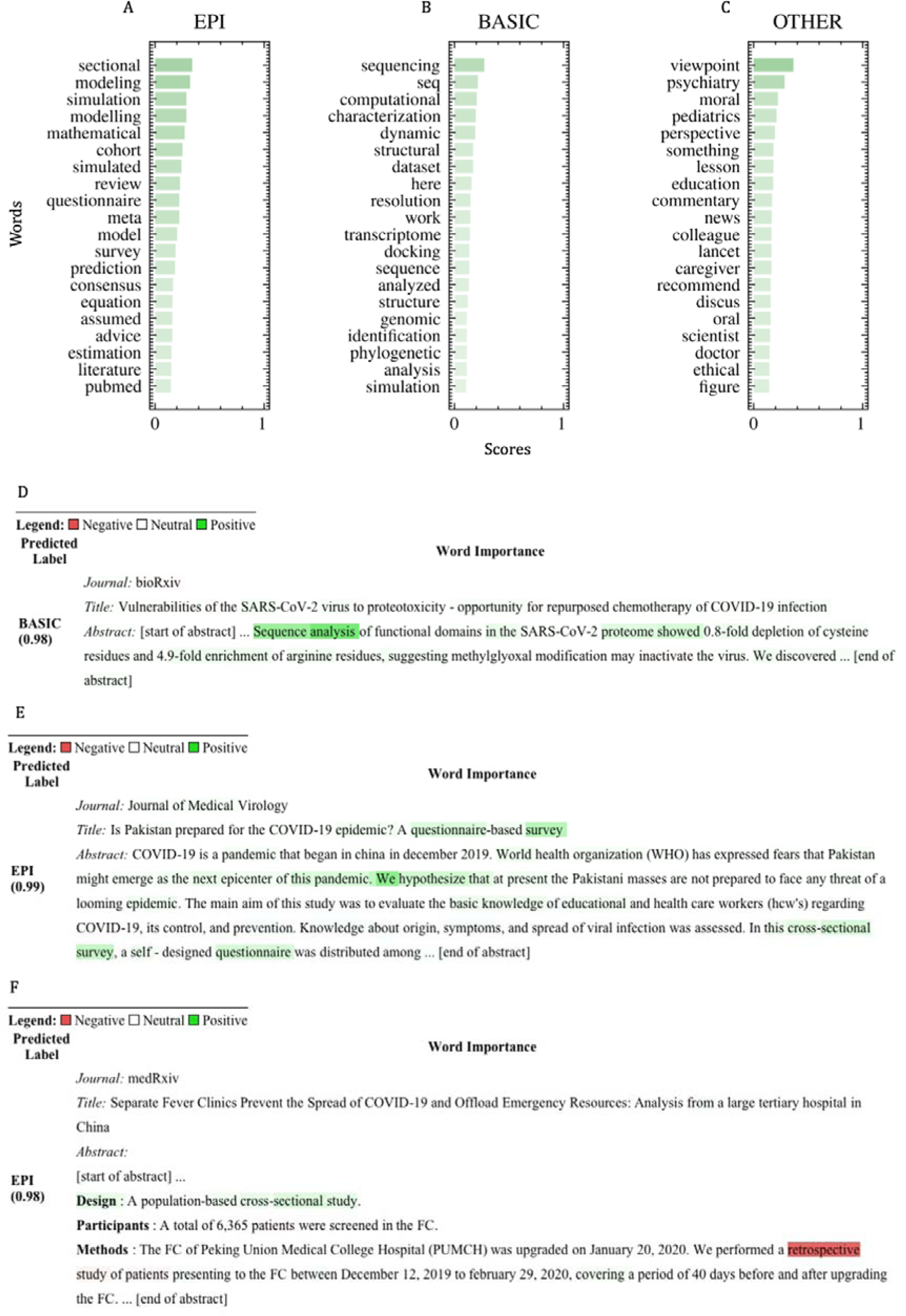
A-B-C: Top 20 positive impact words for either EPI (A), BASIC (B) or OTHER (C) subclasses when taking the integrated gradient on a never-seen set of about 600 documents. D-E-F: Classification examples with a focus on passages with impact words scores.

Figures 6D to 6F depict three publications highlighted using their integrated gradient scores. Publication in Figure 6D^1^ was chosen because it illustrates the usage of the top *BASIC* impact words whereas publications in Figures 6E^2^ and 6F^3^ were selected because they emphasize the highest *EPI* impact words while giving an example of a negative impact word. In Figure 6D, the model predicts the *BASIC* label with 98% probability and the impact words seem to focus on the “sequence analysis” part, with “sequence” being the top impact word in average for that subclass. A look at the sub-subclass prediction level gives a probability of about 95% for the *BASIC: Sequencing and phylogenetics* sub-subclass. In Figure 6E, there is an example of a “sectional” occurrence, the reported most important word for the subclass *EPI*. In our set, the word appears in 7 documents, each time along with the words “cross” and “study”. This publication is classified in *EPI: Cross-sectional study* sub-subclass with a probability of 96%. Interestingly, all 7 documents were classified as *EPI: Cross-sectional study* except for the publication of Figure 6F which was classified as *EPI: Cohort study* with 74% probability and, for which, the classifier seems to give more importance to the word “retrospective” in the methods section than to “sectional” in the design section. As both sub-subclasses are nested into the same subclass, the publication is still classified in the *EPI* subclass with a high probability of 98%.

## Discussion

In this article, we introduce an efficient methodology to assist epidemiologists and biomedical curators to screen articles for inclusion in living systematic reviews by providing a COVID-19 literature triage solution based on deep learning methods. Supported by an existing manually classified collection, we proposed a classification method that automatically assigns categories from a living evidence knowledge base to scientific documents using BERT-like language models, based on which we proposed two methods to combine individual model predictions (probability sum and voting). The results demonstrate that the ensemble performs consistently better than any standalone model, statistically improving upon the best standalone baseline on both strict classification and ranking tasks. Error analyses for the living evidence dataset used in our experiments showed that classification confusion often happens at the intra category level. It helped to explain the difference of performance observed when zooming from sub-subclass to class level, for which micro F1-score goes from almost 70% to almost 90%, respectively. We believe that in this case there are important patterns within categories that the machine learning models can identify and exploit to provide the correct predictions at the class and subclass levels. On the other hand, at the sub-subclass level, we expect that the documents could be often related to more than one category, that is, they are mostly within one category, but may also contain information associated to another category, which could lead to the confusion of the classifier when assigning the sub-subclass, a phenomenon which also occurs during the human annotation. Hence, we believe that a multi-label assignment strategy at the sub-subclass level could be an interesting alternative in the original annotation protocol.

Given the strong performance of the proposed classifier, it could be used to support annotation of scientific articles and help to speed up, augment and scale up epidemiological reviews and biomedical curation. When looking at the problem from a ranking perspective, in which the system suggests a list of sub-subclasses for a given article, the ensemble returned the right category in its top 3 suggestions for almost 90% of the cases. Such a robust performance could help augment the annotation process, for example, by enabling human annotators to double the number of screened articles, replacing an annotator by a machine annotation in the standard double annotation process. In this setting, if the category proposed by the human annotator matched one of the top 3 categories proposed by the automatic classifier, this category would be deemed validated. Otherwise, it would be sent to a senior annotator for a final decision on the remaining 10% of the cases. Considering that a typical inter-annotator agreement in the health and life sciences field is around 80% (47), this setup could reduce the number of human resources required by at least 50% while maintaining the high quality of the annotations. Alternatively, when using a voting strategy with a confidence threshold, we showed that our method was capable of robust and superior performances in a subset of the collection on the class level (about 98.5% F1-score on 50% of the dataset). This approach could be used for example in the triage process, when a large batch of articles needs to be classified, thus scaling up the classification process.

The interpretability analysis showed that the model is not a complete black box as it is often the case in deep learning applications. Using the integrated gradient method helped to understand why the model classified a publication according to a sub-subclass instead of another. These results could be additionally used by annotation experts as a tool to highlight documents during the curation process. It would also be interesting to investigate the results of this analysis at the subclass level, which we believe could lead to a lexicon defining each subclass. Such approaches could then be combined to get multiple views by category level, which could be further assembled to get better publication insights and perhaps better screening results. We leave this investigation for future works.

A main limitation of the study is that it uses a dataset of only one living evidence knowledge base to train and evaluate the models. Thus, it is unclear how the proposed methodology will generalize to corpora and categories used in other reviews and living evidence knowledge bases. That said, given the strong performance obtained in other corpus types by a similar methodology (34), we believe that it shall generalize well. Second, in our experiments, we fail to explore the full contents of the articles. This is due to the unavailability of the full text for a large portion of the collection due to either paywall or restriction by publishers to process full text by NLP pipelines. Additionally, as the time complexity of the models used are quadratic with the number of words, the computation time becomes prohibitive as we move from abstract to full text content. Nevertheless, we believe that valuable information supporting the classification can sometimes only be found in the full text of the manuscripts. An extended version of the approach could investigate such corpora.

## Conclusions

In this work we described an effective methodology to perform automatic classification of COVID-19-related literature to support creation of systematic living reviews and living evidence knowledge bases. The proposed ensemble model provided strong (semi-)automatic classification performance, significantly outperforming standalone methods, and enabled the categorization of a subset of the collection with improved accuracy. Hence, this approach could serve as an alternative assistant to professionals dealing with the COVID-19 pandemic literature outbreak. Ultimately, our method provides a performant and generic procedure, enabling efficient annotation of important volumes of scientific literature, which could be leveraged to assist experts in different literature classification tasks and extended to different types of review methodologies.

## Footnotes

1 https://www.biorxiv.org/content/10.1101/2020.04.07.029488v1.full

2 https://pubmed.ncbi.nlm.nih.gov/32237161/

3 https://www.medrxiv.org/content/10.1101/2020.04.03.20051813v2

## List of abbreviations

BERT: Bidirectional Encoder Representations of Transformers
NLP: Natural Language Processing
COAP: COVID-19 Open Access Project
AUC-ROC: Area Under the Curve of the Receiver Operating Characteristics
CI: Confidence Interval
MAP: Mean Average Precision

## Declarations

### Ethics approval and consent to participate

Not applicable.

### Consent for publication

Not applicable.

### Availability of data and materials

The datasets used and analyzed during the current study are available in the COAP living Evidence database: https://zika.ispm.unibe.ch/assets/data/pub/ncov/. The training, testing and ensemble source codes are available under https://github.com/ds4dh/CovidReview.

### Authors’ contributions

JK designed and implemented the models, ran the experiments and analyses. JK, DT and QH wrote the manuscript draft. NB created the benchmark dataset. DT, PA and NL conceived the experiments. MC, HI and LH programmed and maintained the COVID-19 Open Access Project living evidence database. DBG and AMI organized the annotation of study design in the study records. All authors reviewed and approved the manuscript.

## Acknowledgements

Lucia Araujo-Chaveron, Ingrid Arevalo-Rodriguez, Muge Cevik, Agustín Ciapponi, Muhammad Irfanul Alam, Kaspar Meili, Eric A. Meyerowitz, Nirmala Prajapati, Xueting Qiu, Aaron Richterman, William Gildardo Robles-Rodríguez, Shabnam Thapa, Ivan Zhelyazkov annotated records in the COVID-19 Open Access Project living evidence database.

